# Measuring RBC deformability and its heterogeneity using a fast microfluidic device

**DOI:** 10.1101/2023.07.18.549500

**Authors:** Savita Kumari, Ninad Mehendale, Tanusri Roy, Shamik Sen, Dhrubaditya Mitra, Debjani Paul

## Abstract

We report a high-throughput microfluidic device to determine the Young’s modulus of single red blood cells (RBCs). Our device consists of a single channel opening into a funnel, with a semi-circular obstacle placed at the mouth of the funnel. As a RBC passes the obstacle, it deflects from its original path. Using populations of artificially-stiffened RBCs, we show that the stiffer RBCs deflect more compared to the healthy RBCs. We then generate a calibration curve that maps each RBC trajectory to its Young’s modulus obtained using an atomic force microscope. Finally, we sort a mixed population of RBCs based on their deformability alone. Our device could potentially be further miniaturized to sort and obtain the elastic constants of nanoscale objects, such exosomes, whose shape change is difficult to monitor by optical microscopy.

## INTRODUCTION

Red blood cells (RBCs) are more deformable (i.e, Young’s modulus ∼0.25 - 26 kPa) ^1, 2^ compared to most other cells in the body, which allows them to transport oxygen through the microcirculation. Any change in the deformability of RBCs can have major physiological effects. If RBCs become stiffer, they can occlude the blood flow in capillaries and eventually lead to excessive RBC destruction in the spleen. Healthy RBCs become stiffer when they are stored for several weeks^3^. The deformability of the RBCs changes under certain pathophysiological conditions, such as, malaria^4^, sickle cell disease^5^, diabetes^6^, peripheral vascular disease^7^, etc. This observation suggests that such conditions may be detected by measuring the elastic properties of RBCs. As RBCs are heterogeneous biological objects, we expect their physical parameters to be distributed over a range of values. For example, the variation in the volume of RBCs for a single individual is given by the ‘RBC distribution width’ (RDW). RDW is obtained by dividing the standard deviation in the volume of single RBCs by the mean corpuscular volume of a RBC and multiplying the resulting fraction by 100. It varies from 11% to 15% in healthy individuals^8^. Clearly, we first need to determine the distribution of elastic moduli (e.g. Young’s modulus, shear modulus, etc.) of RBCs in healthy individuals before we can proceed to understand why and how they change in disease. This kind of information can only be obtained from single-cell measurements performed on a large population of RBCs.

There are a number of conventional techniques by which the deformability of RBC populations can be measured at the level of single cells^9^ such as micropipette aspirations^10, 11^, atomic force microscopy (AFM)^12, 13^, membrane fluctuations^14, 15^, optical tweezers^16, 17^, optical stretcher^18^, etc. Most of these techniques require expensive specialized equipment, have a low throughput, and the data acquired by these methods takes a long time to analyze. These challenges led to the exploration of microfluidic techniques^3, 19, 20^. Most microfluidic techniques do not directly measure elastic constants, but rely on accurate monitoring of the shape change of RBCs in real time as they pass through a constriction. The change in shape is often parametrized and the parameters are used as surrogates of elastic coefficients^21^. In one case^19^, the change in shape is compared to the numerical calculation within a model to derive an elastic constant. This approach, while fast, still requires expensive high-speed cameras (2000 - 4000 fps) to track shape changes of cells in real time. Another class of microfluidic techniques measures the pressures required to pass the RBCs through constrictions narrower than their size. The cortical tension of the RBCs is determined from these measurements by using Laplace pressure ^20^.

Based on direct numerical simulations, Zhu et. al.^22^ proposed a microfluidic device, which is somewhat similar to the Rutherford scattering experiment, to separate capsules by their deformability. The main feature of this microfluidic device is a single semi-cylindrical (semi- circular in 2D view) obstacle placed at the mouth of a funnel. In their numerical simulations, capsules with different deformabilities are made to flow towards the obstacle. These capsules deform while going past the obstacle – soft capsules deform more than the stiff ones – and consequently follow different trajectories in the funnel depending on their deformability, eventually exiting the device at different locations. Stiffer capsules deviate more compared to the softer capsules. The purpose of the funnel after the obstacle is to magnify the small differences in the trajectories of the capsules of different deformability as they go past the obstacle. Vesperini, et. al.^23^ experimentally demonstrated the separation of highly polydisperse vesicles based on size and deformability using the same device design. They showed that at low flow rates the capsules are separated based on size (similar to pinch-flow fractionation), while at high flow rates they are separated by deformability. They also introduced a critical capillary number (i.e. the ratio of viscous to elastic forces) between 0.1 and 0.2 to distinguish between these two sorting regimes. In a later report^24^, they demonstrated sorting of capsules of a fixed size based on their capillary number.

We use a similar device design, but go further, by sending red blood cells (RBCs) through this device. Instead of using a high-speed (3000 – 4000 fps) camera to track the shape change of a RBC in real time, we track its path using a 25-30 fps camera. By performing this experiment with a large population of RBCs, we can obtain the distribution of the tracks of the entire RBC population. In parallel, we also perform AFM measurements on a population of healthy RBCs collected from the same healthy individuals to obtain a distribution of their Young’s moduli. We then compare the distribution of RBC trajectories with the distribution of AFM results and minimise their Jensen-Shannon divergence. In the process, we generate a calibration curve for this device that can uniquely map each trajectory in the microfluidic device to a corresponding Young’s modulus for the RBCs.

This technique of measuring the deformability of RBCs is both cheaper and simpler compared to the other microfluidic methods as we do not need to fabricate devices with very small feature sizes (i.e. comparable to the lateral dimension of RBCs, which is ∼ 2 *μ*m or less) or use very high-speed (i.e. several thousands of fps) cameras. Using our calibration curve, we can also accurately determine the distribution of the Young’s modulus of a mixed population of RBCs consisting of deformable and stiff cells. Furthermore, the observation that healthy and stiff RBCs exit the device at different angles suggests the potential of our device for sorting of cells based on their mechanical properties. The device can, in principle, be scaled down by a factor of hundred to measure and detect the deformability of extracellular vesicles, which are so small that they cannot be resolved under the optical microscope.

## RESULTS AND DISCUSSION

### RBCs of different deformability follow different trajectories in our microfluidic device

We fabricated a microfluidic device design (**fig. 1A**) that is similar to that numerically proposed by Zhu et al^22^ and reported by others for measuring the deformability of spherical capsules^24^. It has three functional sections for (a) flow focusing (see also supplementary fig. **S1**), (b) deformability measurement of single RBCs, and (c) collection of the sample fractions with different deformabilities. As shown in **fig. 1B**, guided by the work of Zhu et al, we hypothesize that the softer RBCs will be deflected from their paths by a smaller angle, while stiffer RBCs will be deflected by larger angles. We first flow a dilute solution of healthy RBCs along a short and straight channel segment towards the obstacle. After crossing the obstacle, different RBCs follow different trajectories and exit the device at different angles. We record the trajectories of these RBCs as they move along the funnel after passing the obstacle.

**Figure 1.**
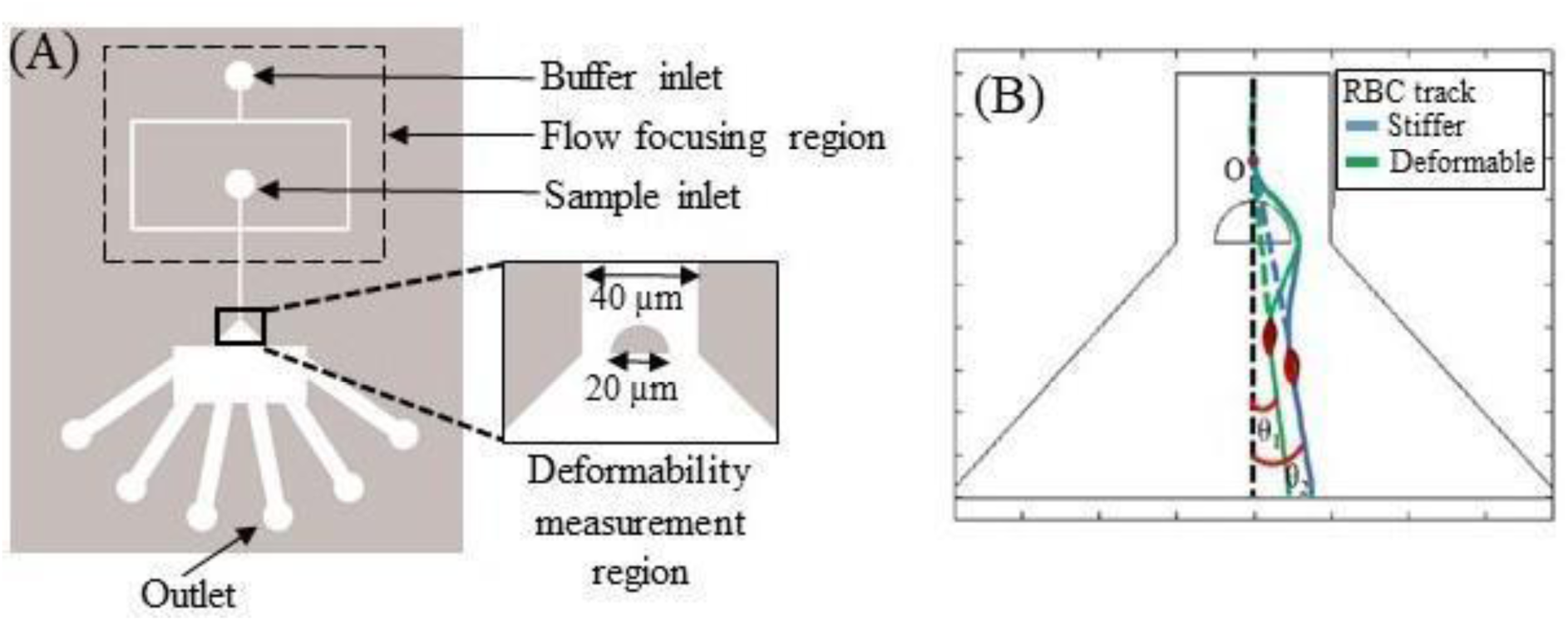
Schematic diagram of the microfluidic device to deflect healthy and stiff RBCs from their paths. (A) The device has a 40 μm-wide straight channel that opens into a funnel. A semi-circular obstacle of 20 μm diameter is placed at the mouth of the funnel. A flow-focused and very dilute stream of RBCs moves along the mid-plane of the straight channel towards the obstacle, allowing only one RBC to approach the obstacle at a time. (B) Softer RBCs are deflected by a smaller angle θ_1_, while stiffer RBCs are deflected by a larger angle θ_2_ in this device geometry.

We also perform a COMSOL simulation to solve the Navier-Stokes (not Stokes as our Reynolds number is about 10) equations for the background flow in absence of RBCs in the device to obtain the streamlines of the flow. These streamlines are numbered from 0 to 50, starting from the central streamline. We then superpose experimental images of RBCs in our device with the streamlines obtained from COMSOL simulation to map the centre of mass of each RBC in the funnel to a unique streamline. Finally, we count the number of the RBCs moving along each streamline and plot the data as a histogram in **fig. 2** (top panel).

**Figure 2.**
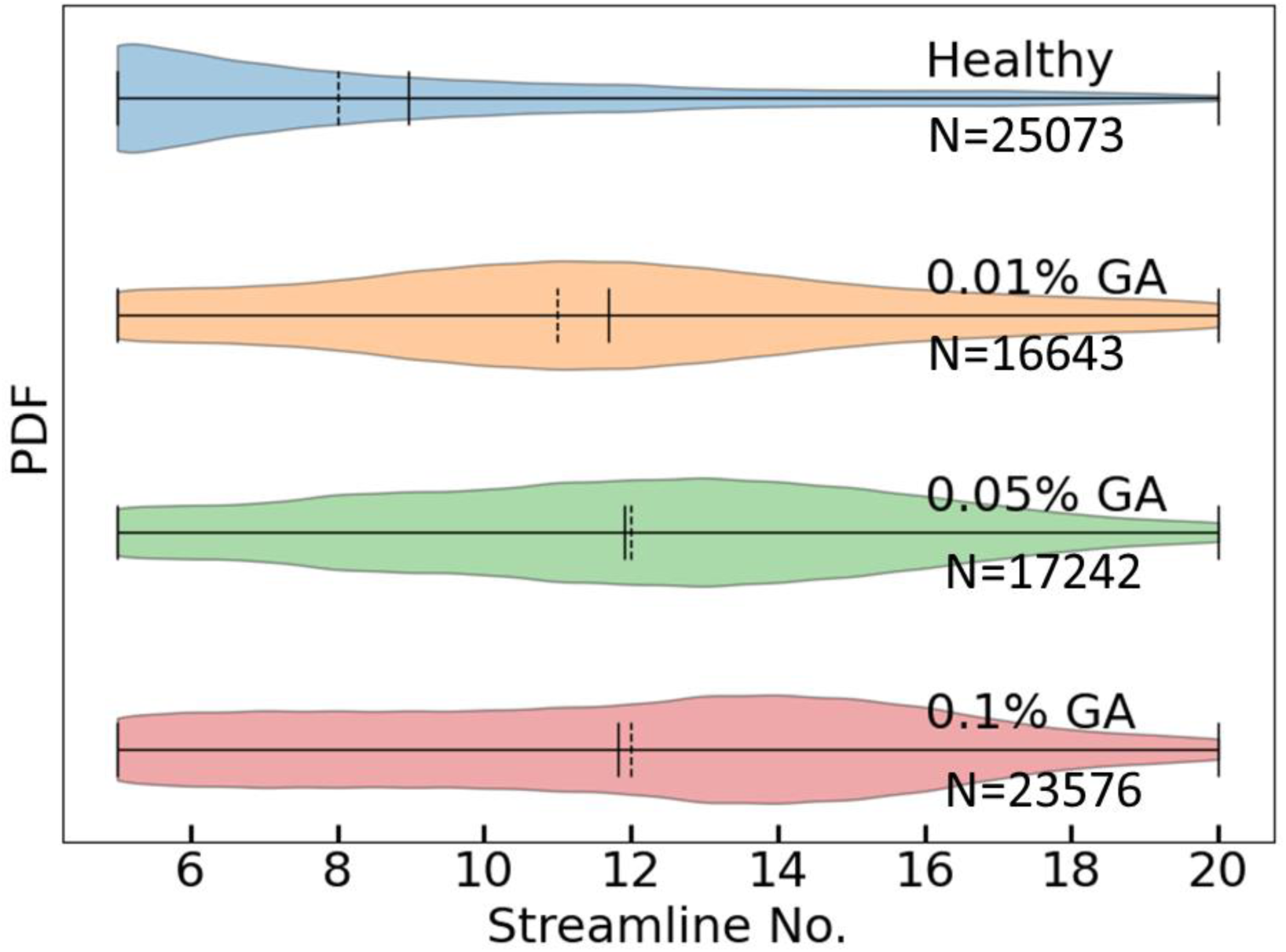
Probability distribution function showing the tracks of different RBC populations along different streamlines in the microfluidic device. Each panel denotes a RBC population treated with a specific concentration of a chemical stiffening agent glutaraldehyde (GA). N is the number of RBC tracks analysed to obtain a particular distribution. Dashed vertical lines indicate the median and continuous vertical lines indicate the mean of the distribution. Untreated RBCs follow the streamlines with lower numbers, while the progressively stiffer RBCs are found along the streamlines with higher numbers.

Next, we use several batches of RBCs – we treat each batch with a specific concentration (0.01%, 0.05% and 0.1% respectively) of a chemical stiffening agent glutaraldehyde (GA) to generate artificially stiffened populations of RBCs with different mean stiffness values – and repeat the same experiment. GA is widely used to attach RBCs to different surfaces and also to reduce their deformability^25^. The successive panels in **fig. 2** show the probability distribution functions (PDF) of positions of the RBCs along different streamlines when these are treated with different concentrations of GA. We also measure the diameters of untreated and 0.1% GA-treated RBCs to ensure that GA-treatment does not change the sizes of the RBCs, and hence, the resulting RBC tracks in the funnel region is only due to the changes in their deformability. In the next section we show how we use this data to obtain the deformability distribution of the RBC populations.

### Measurement of the deformability of single RBCs

At this point, recall what we understand by the ‘deformability’ of a single RBC. As the simplest example, consider a three-dimensional, homogeneous and isotropic material. For small deformations, a complete description of the elastic properties of this material is given by two scalars (for example, the two Lamé coefficients). Since RBC is a complex biological object, it is not obvious how many independent scalars are necessary to completely describe its elastic properties. Measurements of the elastic properties of RBC have a long history of using different methods, e.g., AFM, micropipette aspiration, optical tweezers, membrane fluctuations, etc. Each technique may end up measuring a different aspect of the deformability of RBCs. For example, AFM measurements yield an effective three-dimensional Young’s modulus for RBCs, while micropipette aspiration measures the bending modulus. Hence, it is not possible to directly compare the results of these techniques with each other unless different methods are cross-calibrated against each other using the same RBC^26^. Finally, the results of a particular class of measurements are interpreted using several different models of the RBC, which makes the measured constants model-dependent. One class of measurements attributes the results completely to the elasticity of the RBC membrane, while ignoring the inner constituents of the cell. These techniques measure different elastic constants of the membrane: (a) shear modulus (measured by micropipette aspiration^10^, optical tweezers^16^, magnetic twisting cytometry^26^, and membrane fluctuations^14^), (b) bending modulus (given by micropipette aspiration^27^ and membrane fluctuations^14^), (c) area extension modulus measured using micropipette aspiration^28^, and (d) the aspect ratio^29^ of the RBC, which is not an elastic modulus but can be used as a somewhat indirect measure of deformability. Another class of measurements attributes their results to the entire RBC as a solid object. They interpret their results in terms of the Young’s modulus measured using an AFM^30, 31^. Finally, there is a class of experiments^32, 33^ that reports the transit times or the velocities of soft or stiff RBCs through a constriction as surrogate measures of their deformability. Since these measurements do not yield an actual elastic modulus, we will not discuss those in this paper.

We need a calibration curve to assign a unique value of an elastic modulus to each streamline in our device. To construct the calibration curve, it would have been ideal if we could design capsules with the same shape as an RBC, but with a membrane whose elastic constant is known by other measurements. As this is impossible at this point, we must resort to one of the conventional methods of RBC deformability measurement. Therefore, we perform atomic force microscopy (AFM) on a population of RBCs collected from the same healthy individuals whose blood samples were used for the microfluidic measurement. These measurements allow us to obtain a PDF of Young’s moduli of healthy RBCs, as shown in **fig. 3**.

**Figure 3.**
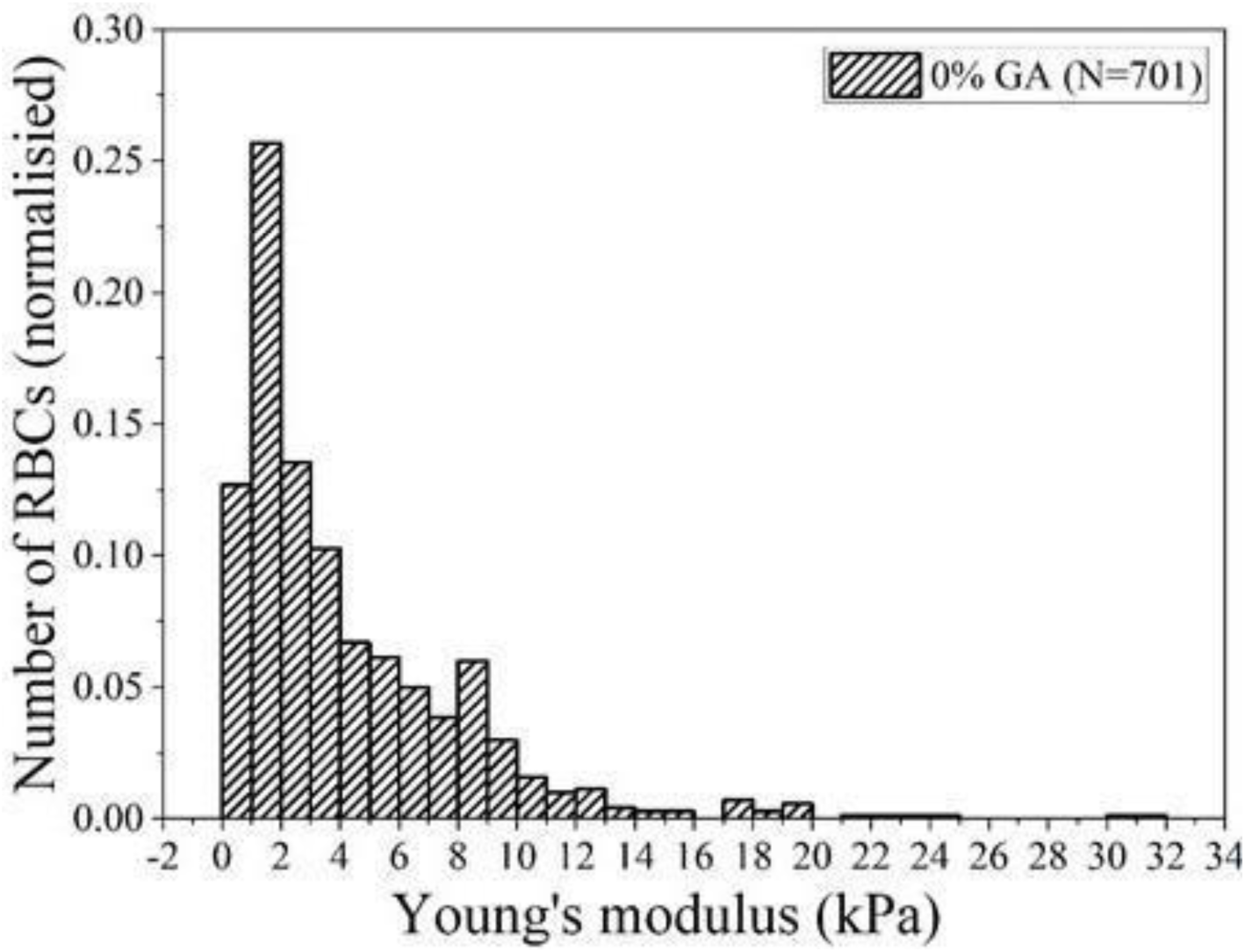
Probability density function (PDF) of Young’s modulus of RBCs (N = 701) obtained by AFM measurements. These RBCs were obtained from the same healthy individuals whose blood samples were used for microfluidic measurements. N indicates the number of RBCs.

A calibration curve for a RBC in the Rutherford device is given by the function Y = f(s) where Y indicates the Young’s modulus and s indicates the streamline number. In addition to Young’s modulus of RBC, the calibration curve must also depend on the flow rate, the exact geometry of the device, the viscosity of the fluid, and the initial undeformed shape of the RBC. If all these parameters, except the Young’s modulus, remain the same, then the calibration curve is a one-to-one map from the streamline number to the Young’s modulus. The best possible way to find this calibration curve would be to send a single RBC through the Rutherford device, measure the streamline it comes out along and then measure the Young’s modulus of this RBC in an AFM. This is impossible to perform due to logistical reasons. Hence, we take a data-based approach.

Let P(Y) be the probability distribution function of the Young’s modulus of a family of RBCs and p(s) be the probability distribution function of the streamline of the same family of RBCs. Let us assume f to be the calibration curve function such that

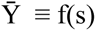

and calculate Q, which is defined as the PDF of Ȳ. If we knew the calibration curve (f) precisely, then we would obtain P (Y) = Q (Ȳ). We try to find the best calibration curve our measurements allow. Thus, the calibration curve is the function f(s) that minimizes the distance between the two *experimentally measured* PDFs P and Q. As a measure of the distance we use the Jensen-Shannon divergence:

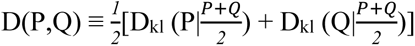

Here D_kl_ is the Kullbac--Leibler divergence defined by:

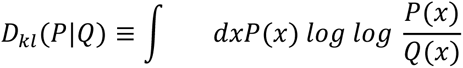

Note that D_kl_ (Q|P) ≠ D_kl_ (P|Q).

But D(P,Q) is symmetric in P and Q. It can be considered as a squared metric^34^. If P = Q then D(P,Q) = 0.

To perform the minimisation of the Jensen--Shanon divergence we write the calibration curve as a polynomial

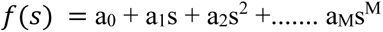

The minimisation problem then reduces to a problem in M dimensional space.

Furthermore, we impose two constraints: The calibration curve must be monotonic i.e, all the a_k_ values must be positive. Also, it must be differentiable. This condition is satisfied by choosing a polynomial. Furthermore, we limit each a_k_ within 0 and a_k_^max^ with the maximum value chosen such that

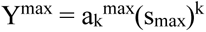

where Y^max^ and s_max_ are the maximum measured values of the Young’s modulus and the streamline number, respectively. The minimisation is done using the particle swarm algorithm. The constraints can be imposed in a straightforward manner by putting boundary conditions on the domain over which we search for the minimum. We limit ourselves to M=4. Clearly, for s=0, Y=0, which implies a_0_ = 0. The calibration curve thus obtained is shown in **fig. 4**. It maps each streamline to a single value of young’s modulus (Y).

**Figure 4.**
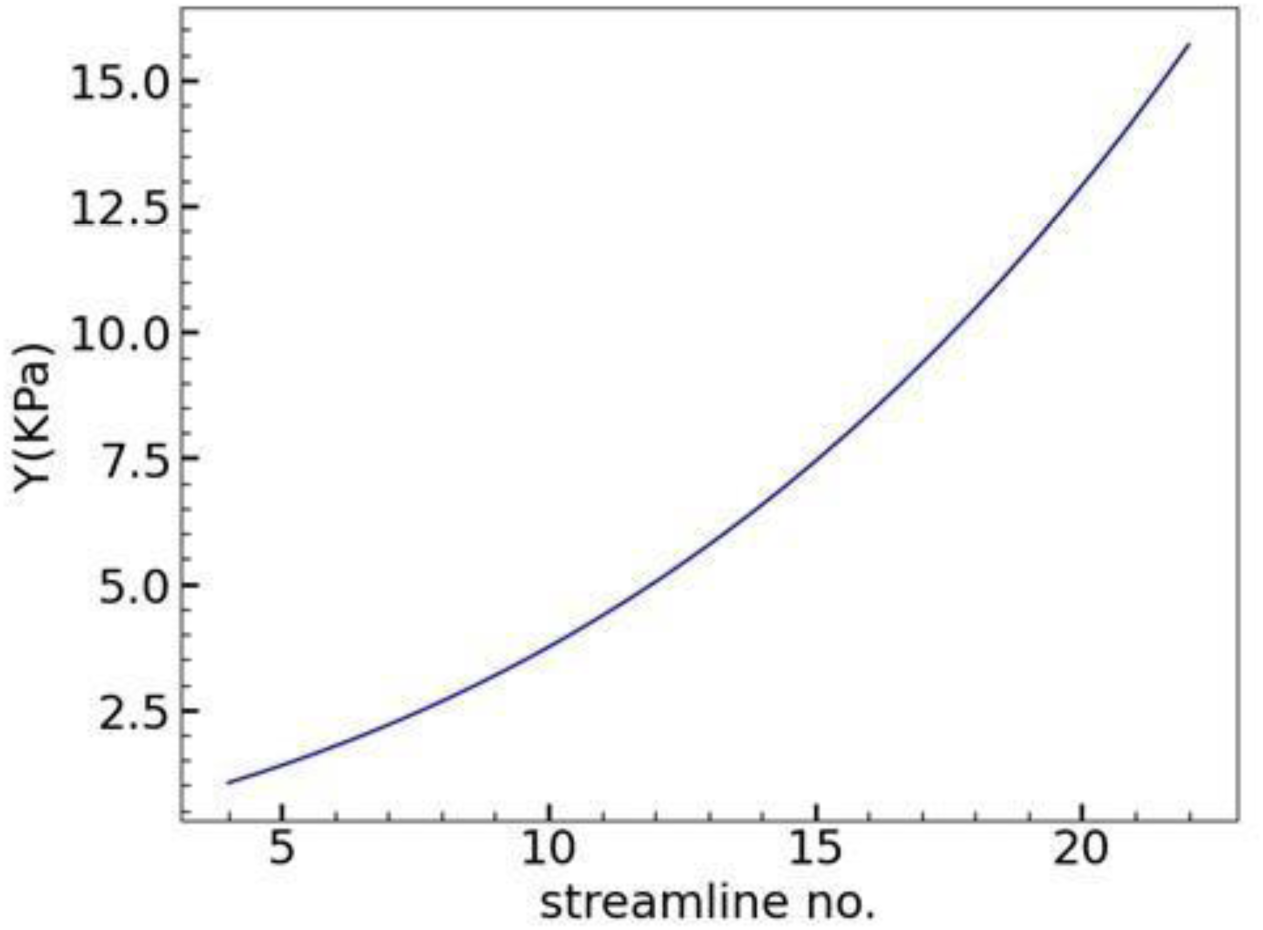
Calibration curve obtained by comparing the PDF of RBC trajectories with the PDF of AFM results. It maps each RBC trajectory to a unique value of Young’s modulus for the corresponding RBC.

From our experiment, we can thus generate a PDF of Young’s modulus for healthy and artificially stiffened RBCs, as shown in **fig. 5**. In this plot, the shaded region shows the probability density function. The Young’s moduli values for healthy and artificially stiffened RBCs are 3.6 ± 0.02 kPa (mean ± SEM) (healthy or untreated), 5.26 ± 0.02 kPa (treated by 0.01% GA), 5.41 ± 0.02 kPa (treated by 0.05% GA) and 5.38 ± 0.02 kPa (treated by 0.1% GA). The mean Young’s modulus obtained by our measurements matches with the value reported in the literature^1, 13, 31, 35^. While GA is widely used in biology to stiffen RBCs and attach them to surfaces^2, 30^, at the moment there is no reported systematic characterization of the elastic constants of RBCs after treatment with specific concentrations of GA. Our results provide a way to controllably characterise the deformability of RBC populations after treating these with different GA concentrations.

**Figure 5.**
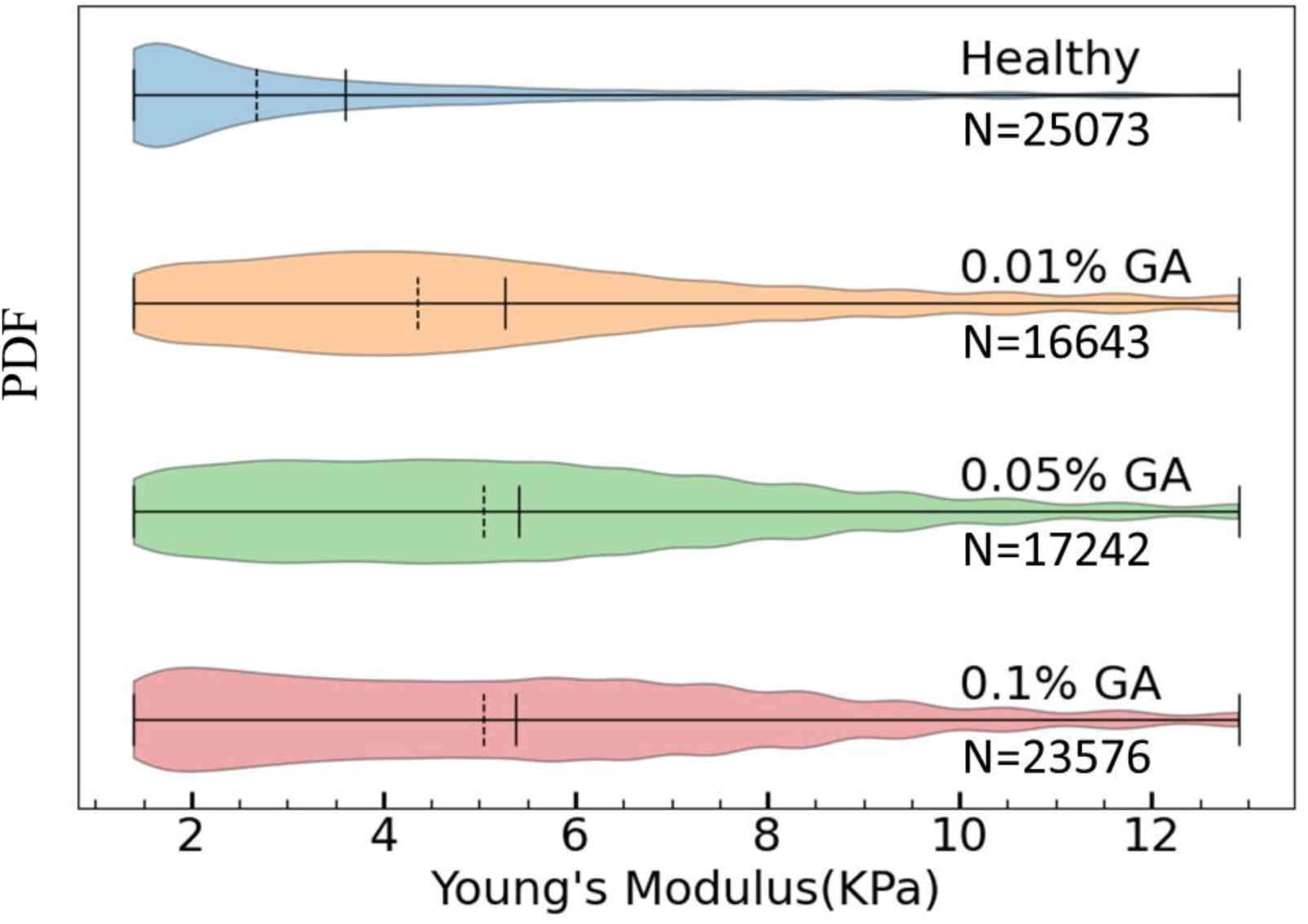
PDF of Young’s modulus of healthy and chemically stiffened RBCs obtained using our calibration curve. The RBCs are stiffened by treating them with different concentrations of glutaraldehyde (GA) varying from 0.01% to 0.1%. The vertical solid line shows the mean and dashed vertical line shows the median of the distribution. The mean Young’s moduli values for healthy and artificially stiffened RBCs are 3.6 kPa (healthy or untreated), 5.26 kPa (treated by 0.01% GA), 5.41 kPa (treated by 0.05% GA) and 5.38 kPa (treated by 0.1% GA).

### Sorting of RBCs from a mixed population based on their deformability

Finally, we check the ability of our device to measure deformability of single RBCs from a heterogeneous mixture of 50% healthy RBCs and 50% chemically stiffened (with 0.1% GA) RBCs. We pass this mixed population of RBCs through our device. We track their trajectories as they move through the device and calculate the PDF for Young’s modulus by using the calibration curve. As shown in **fig. 6**, we obtain two peaks in the PDF for the mixed population: one corresponding to healthy RBCs at 2.5 kPa and other corresponding to stiffened RBCs at 6.4 kPa. These results suggest a potential use of this device for sorting RBC populations based on their deformability alone.

**Figure 6.**
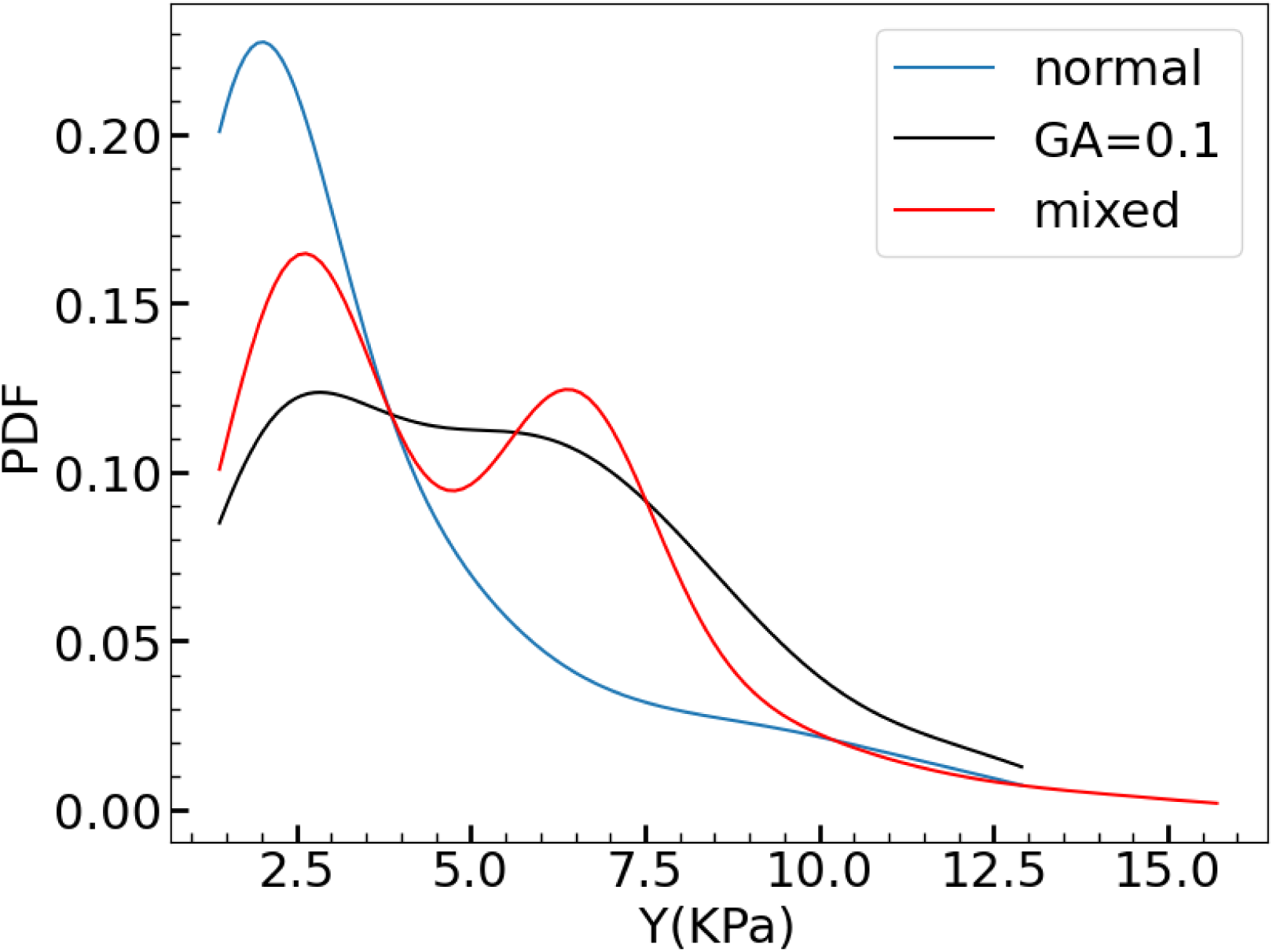
PDFs of a population of healthy RBCs (N = 25073), a population of chemically stiffened RBCs (N = 23576) and a mixed population of RBCs (N = 36388) obtained by mixing 50% healthy and 50% chemically stiffened (0.1% GA) RBCs. N indicates the number of tracks that were analysed to obtain the distribution.

In summary, we report a microfluidic device to obtain the distribution of the Young’s modulus (Y) of healthy and chemically stiffened RBC populations. The mean Young’s modulus of healthy RBCs obtained by our method is in the same range as those obtained by AFM in the reported literature. We also pass a mixed sample containing equal numbers of healthy and stiff RBCs through our device, and demonstrate that the PDF of Young’s modulus has a clear bimodal distribution, with peaks at very similar locations compared to the ones obtained with individual populations. Thus, we can use our device to separate populations of RBCs based on deformability. Furthermore, our microfluidic device does not need fabrication of any feature size smaller than 10 *μ*m. It can work with a regular 25 - 30 fps microscope camera to measure ∼ 25,000 RBCs in each experiment at an average rate of 10 cells/s. Therefore, this method can be an inexpensive and fast alternative to the other deformability cytometry technologies currently available for measurement of elastic constants of red blood cells. Further, it is ideally suited to construct a probability distribution function of deformabilities of a family of RBCs. Finally, our techniques can potentially be used to both measure and sort nanometer-sized objects such as *extracellular vesicles* by size and deformability. These are about a hundred times smaller than RBCs and lie at the limit of resolution of optical microscopes. They can be observed as bright point sources under high resolution microscopes, but their shapes under mechanical stress cannot be imaged by the existing techniques. So conventional deformability-based cytometry^19^ cannot be used to measure their elastic moduli. As our method relies on measuring the trajectory of the elastic object, and not its shape, it is more easily generalizable to such nanoscale objects.

## EXPERIMENTAL PROCEDURES

### Resource Availability

#### Lead Contact

Further information and requests for resources and reagents should be directed to and will be fulfilled by the Lead Contacts, Dhrubaditya Mitra or Debjani Paul.

#### Materials Availability

All materials are available on request from the authors.

#### Data and Code Availability

Data is available on request from the authors.

#### Design and fabrication of the microfluidic device

The microfluidic device has three sections. The first section focuses a dilute stream of RBCs to ensure that they move along the centre of the channel. The second section consists of a 40 µm-wide straight channel opening into a 45° funnel, with a single semi-cylindrical (semi- circular in 2D view) pillar of 20 µm diameter placed at the junction of the straight channel and the funnel. The flat side of the pillar faces the funnel, while the curved side faces the focused stream of RBCs. The third section has six outlets, separated from each other by 18° at the end of the funnel to collect the cell fractions with different deformabilities. Our design has a few minor differences from the designs reported earlier^23, 24^. First, the gap between the obstacle and the channel wall is 10 µm in our device, which is larger than the diameter (∼ 7 µm) of a typical RBC. This gap is much smaller than the gap (∼ 50 μm) reported for separating spherical capsules^24^. Second, our device height is 5 µm throughout to ensure that the biconcave RBCs approach the pillar in a flat orientation instead of flipping sideways. Allowing for a larger height leads to flipping of the RBCs, which is avoided in our design. Third, there are six outlets instead of five for collecting different sample fractions with different deformabilities.

The devices are made from the elastomer PDMS (Sylgard 184 from Dow Corning, India) by soft lithography^36^. We follow our reported protocol^37^ to make SU-8 masters for lithography with a height of 5 μm. We then mix the two parts of PDMS in a 10:1 ratio, pour it on the master and cure in an oven at 65°C for 45 min. We punch inlets and outlets in the cured PDMS chip using a 1.5 mm diameter biopsy punch (Accu-Sharp from Med Morphosis LLP, India), and bond the chips to a 24 mm x 40 mm x 0.13 mm cover slips (Blue Star, India) using oxygen plasma (PDC 32G from Harrick Plasma, USA) for 2 min. We then incubate the bonded device in an oven at 65°C for 30 min. The PDMS devices are used without any surface treatment.

### Preparation of blood sample

The experimental protocol was approved by the Institute Ethics Committee of Indian Institute of Technology Bombay (approval number IITB/IEC/2017/020). Seven (four female and three male) healthy volunteers in the age group of 25 - 35 years participated in the study. We collect a drop of blood obtained by a finger prick from each healthy volunteer after obtaining informed consent. We prepare 28% OptiPrep (Sigma Aldrich, India) solution using normal saline and dilute the blood by 500X - 1000X using this solution to avoid settling of RBCs in the syringe or tube during the experiment. To obtain chemically stiffened RBCs, we prepare glutaraldehyde (GA) solutions in 28% OptiPrep and dilute RBCs by 500X - 1000X in it. We treat blood with 0.01%, 0.05% and 0.1% concentrations of GA for 20 min at room temperature to stiffen them to different extents. To measure the tracks of mixed samples in our device, we first prepare 2.5 ml of 0.1% GA-treated blood and mix it with 2.5 ml of untreated blood to obtain 5 ml of the mixed sample.

### Deformability cytometry of RBCs in the microfluidic chip

We use 500X - 1000X diluted blood for deformability measurement. Since the blood sample is heavily diluted and our device height is 5μm, the presence of WBCs or platelets do not affect our results. RBCs suspended in the OptiPrep solution are passed through the sample inlet, whereas 0.9% normal saline is flown through the buffer inlet for focussing the sample stream. The focussed stream of RBCs is allowed to flow towards the obstacle at a flow rate of 1 µl/min. After passing the obstacle, the RBCs leave the device through one of the six outlets. To track the trajectories of the RBCs near the obstacle, we image the RBC tracks using a digital inverted microscope (Cilika from MedPrime Technologies, India) fitted with a 40X (0.6 NA) objective lens. The videos of RBC tracks are acquired at a resolution of 720 x 1280 pixels in uncompressed format by a 30 fps iPad camera fitted to the microscope, while keeping the obstacle at the left-centre of the frame.

### COMSOL simulation

We use the built-in microfluidics module (single-phase laminar flow) of the COMSOL multiphysics software (version 5.2) with ‘no slip’ boundary conditions and a fine mesh. The simulations are performed with a flow rate of 1 μl/min to match the experimental conditions.. To save memory, we simulate the flow in 80 μm length of the straight channel and 40 μm length of the funnel. To label the tracks of the RBCs, we numerically solve the background flow (i.e. the flow without any RBC) with a grid resolution that is consistent with the resolution of our camera (0.9 μm). We number the streamlines obtained from our simulation from 0 at the centre of the funnel to 50 on each side.

### Mapping RBC tracks to streamlines

We extract all frames of the video and use a MATLAB image processing algorithm to obtain the cell tracks. We read and store all frames as a 4D matrix. The number of the frames and their spatial resolution are then determined. We convert each frame from RGB to grayscale and store the first frame of each video (**fig. 7A**) separately. To overlap it with the streamlines obtained from the COMSOL simulation (**fig. 7B**) of the background flow, we calculate the alignment operation (e.g. rotation, scaling and translation parameters (**fig. 7C** and **7D**). As indicated by the three red dots on the **fig. 7A**, we use three coordinates on the obstacle to calculate the alignment factors. (*x*_1_, *y*_1_) and (*x*_2_, *y*_2_) are the coordinates of the two corners of the obstacle respectively. Along with these two coordinates, we use the coordinate (*x*_3_, *y*_3_) of the centre of the curved face to check the orientation of the obstacle.

**Figure 7.**
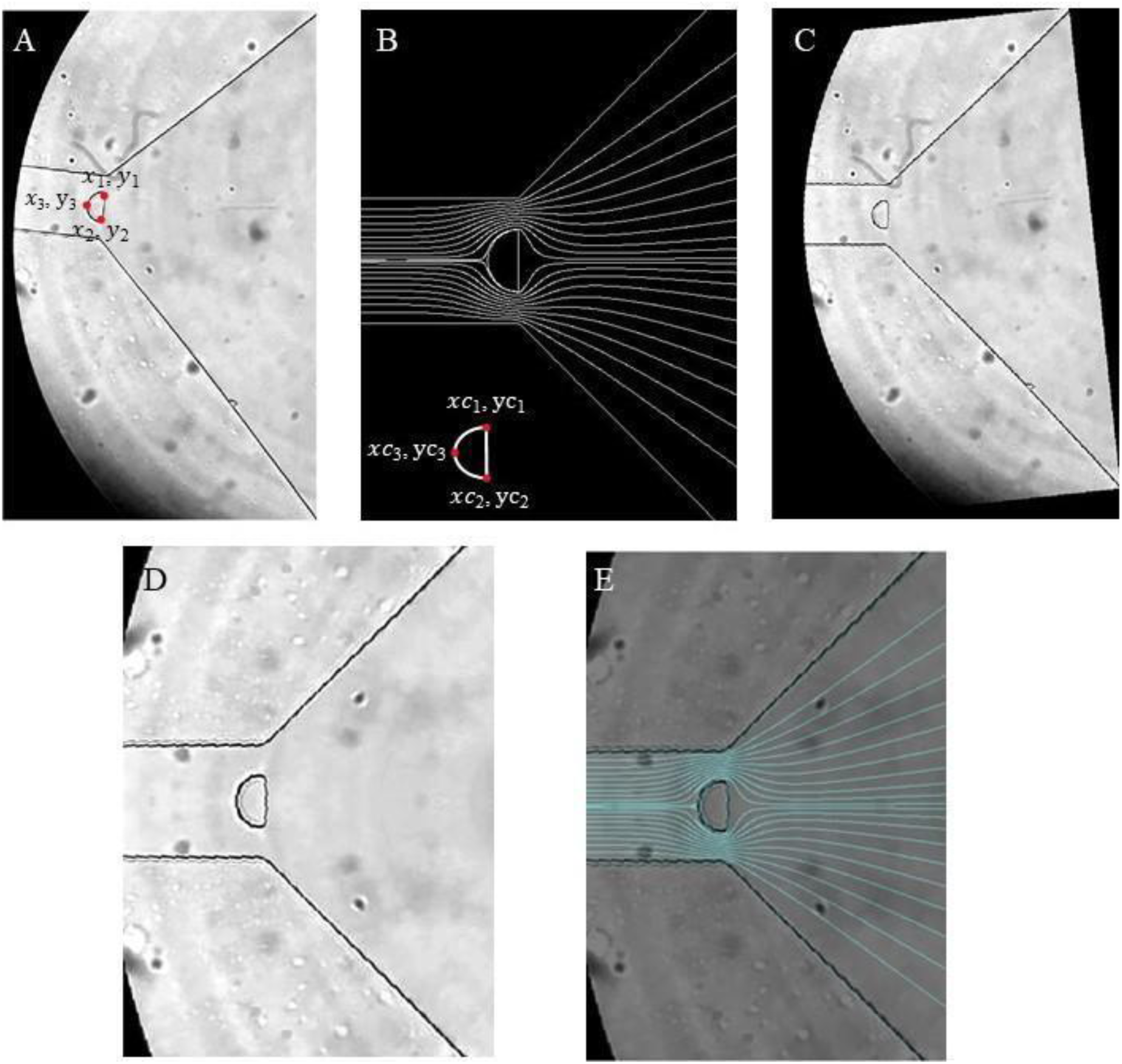
Overlaying an image frame from the experiment to the streamlines obtained from the simulation. (A) and (B) indicate the image frame from the experiment and the frame from COMSOL simulation showing the streamlines, respectively. (C) By comparing the coordinates of three points on the obstacle (shown by red dots), the image frame is rotated by an angle 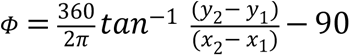. (D) Subsequently, the image frame is scaled by a scale factor of 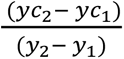 and then cropped. (E) The image shows overlapping of the experiment frame with the streamlines obtained from COMSOL simulation to map the centre of mass of each RBC to a streamline.

Equations (1) and (2) show the formulae that we use for image rotation, where (*x*, *y*) are the coordinates of the pixel to be rotated and (*xCosϕ* − *ySinϕ*, *xSinϕ* + *ySinϕ*) are the new coordinates of the same pixel after rotation by an angle *ϕ*.

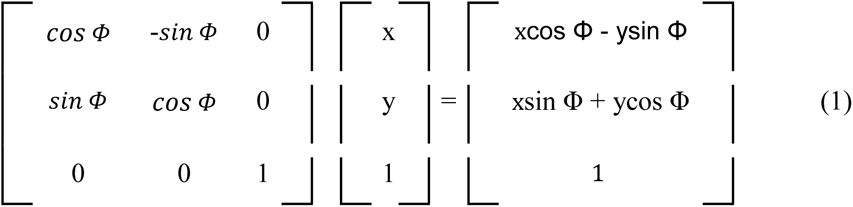

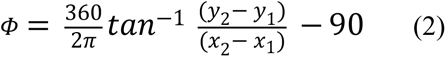

The rotation angle Φ is given by equation (2). If x_3_ > x_2_, a rotation angle (*ϕ* + 180) is used. Equation (3) gives the scale factor needed to resize the image so that it fits on the COMSOL simulation window. The scale factor (SF) is the ratio of the distance (*yc*_2_ − *yc*_1_) between the two corners of the obstacle in the COMSOL simulation to the distance (*y*_2_ − *y*_1_) between the same two points in the experimental image frame.

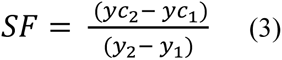

After rotating and scaling the image, we match the coordinates (*xc*_1_, *yc*_1_) in the COMSOL image to the corresponding point (*x*_1_, *y*_1_) in the video frame by a translation operation. Finally, we crop the image and overlay it with the COMSOL simulation window as shown in **fig. 7E**.

To generate the RBC trajectory, we subtract two consecutive frames. This difference gives us the overlapping areas of a RBC at two different time points. The difference image is then converted from grayscale to a binary image using a threshold value of 0.05. We then check whether the feature in the difference image thus obtained is indeed a RBC by measuring its area. It is taken to be a RBC if its area is more than 50 pixels (corresponding to 6 µm diameter).

The streak is then filled using morphological operations and its centroid is determined. The position of the centroid is next multiplied with all possible streamline patterns generated by COMSOL. The only streamline corresponding to a non-zero product after multiplication is chosen as the correct streamline indicating the track of the RBC.

We ignore any tracks mapped to streamlines 1 to 5 as the radius of the objects following these streamlines is too small (∼ 1 μm) to be RBCs. Similarly, all streamlines higher than 23 correspond to objects whose sizes are too large (with radius ∼ 4.6 μm and higher) to be RBCs. We then divide the entire range between streamlines 6 and 22 into several bins. The bin size is set by the resolution (0.9 μm) of our image acquisition system.

### Attaching RBCs to a glass surface for atomic force microscopy

We clean 18 mm diameter glass coverslips in oxygen plasma for 2 min. We then immerse the cover slips in 0.1% (w/v) poly-l-lysine (Sigma Aldrich, India) solution for 30 min to promote adhesion of RBCs to the glass surface. After washing the coverslips with deionized water to remove excess poly-l-lysine, we dry them under vacuum for 2 h at room temperature. We add 100 µl of diluted blood on the treated glass coverslip and incubate at room temperature for 30 min. We wash the coverslip with normal saline to remove any unattached RBCs and immediately perform atomic force microscopy (AFM) measurements.

### AFM measurements

We measure the Young’s moduli of RBCs using a bio-AFM (Asylum Research, USA) in contact mode with a liquid-cell set-up. We use a 10 kHz stiff pyramidal silicon nitride probe (Oxford Instruments, UK) cantilever, with a spring constant of ∼ 0.03 N/m and a half angle of 35°, to measure the deformability of single RBCs. All measurements are done in 0.9% normal saline at room temperature. We measure the force-indentation curve of at least 50 RBCs for each blood sample. We then fit the Hertz model to the first 500 nm of the experimental force-indentation curves to determine the Young’s modulus of each RBC.^38^

## ACKNOWLEDGEMENTS

The authors thank the nanofabrication facility (IITBNF) and the bio-AFM facility at the Indian Institute of Technology Bombay. The authors also acknowledge Vijay Mistari for help with AFM. D.P. acknowledges partial financial support from a seed grant (RD/0512- IRCCSH0-024) from the Indian Institute of Technology Bombay. D.M. acknowledges the support of the Swedish Research Council grant number 638-2013-9243 as well as 2016- 05225. D.M. thanks Apurba Dev for illuminating discussions.

## AUTHOR CONTRIBUTIONS

S.K., N.M., D.M. and D.P. designed the research. S.K. and N.M. performed the experiments and analyzed the data. S.K., N.M. and D.M. performed the COMSOL simulations. S.K., T.R. and S.S. performed AFM experiments and analyzed the data. S.K., N.M., D.M. and D.P. developed the results and wrote the paper. D.M. designed the optimization problem.

## SUPPLEMENTAL INFORMATION

### 1. Effect of flow focussing

**Figure S1:**
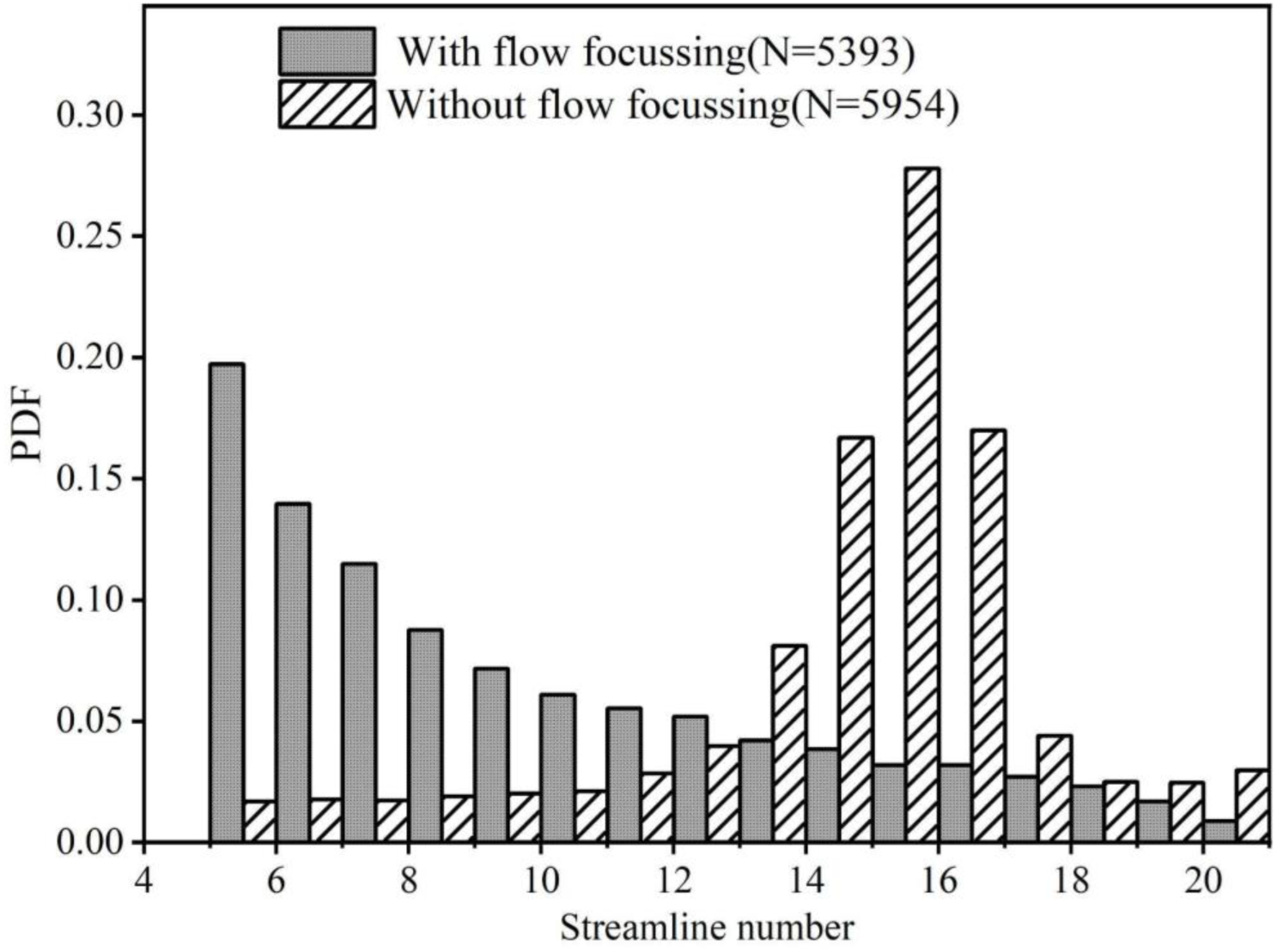
Probability distribution function of the RBC tracks after passing the obstacle, when the RBCs are flow-focussed (hatched pattern) and when there is no flow focussing (solid pattern). We note that in absence of flow focussing, healthy RBCs tracks lie on the entire range of streamlines. Flow focussing ensures that the RBCs approach the obstacle along the channel axis and any deviation in the path is due to its interaction with the obstacle.

### 2. Comparison of RBC sizes with and without glutaraldehyde treatment

**Figure S2.**
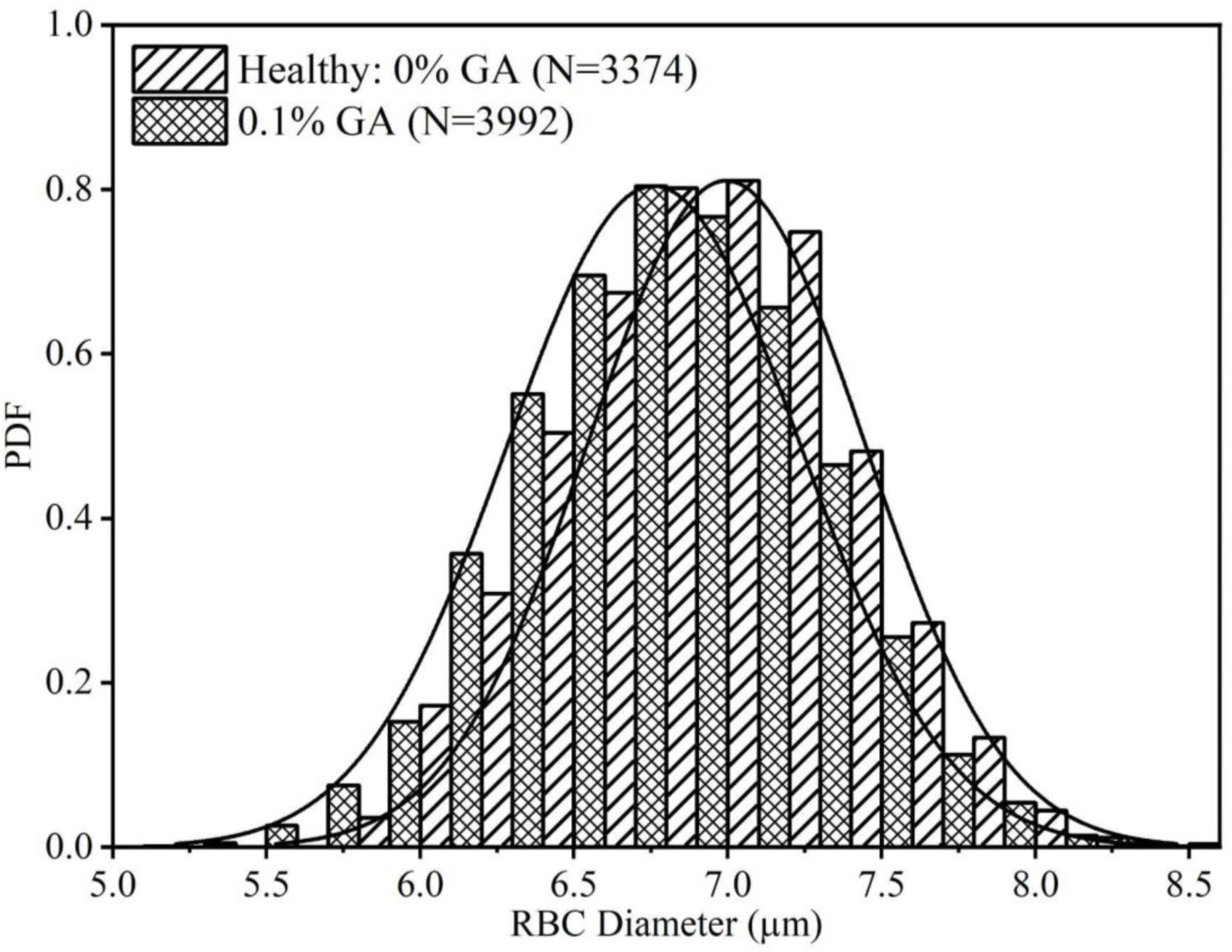
Probability distribution function (PDF) of diameters of healthy (untreated) RBCs and RBCs treated with 0.1% glutaraldehyde. N indicates the number of RBCs. The solid lines indicate normal distributions fitted to the data points. We note that the two PDFs have significant overlap, indicating that the RBC sizes do not change much due to the glutaraldehyde treatment. This implies that the RBC tracks in our device are due to their deformability and not their size variation.

### 3. Movie captions

Movie M1: A time lapse video of the RBC tracks after they encounter the obstacle.The video was acquired at 30 frames per second using the iPad camera of the digital inverted microscope.

